# A *Drosophila* model of mitochondrial disease phenotypic heterogeneity

**DOI:** 10.1101/2023.12.05.570102

**Authors:** Lucy Granat, Debbra Y. Knorr, Daniel C. Ranson, Ram Prosad Chakrabarty, Navdeep S. Chandel, Joseph M. Bateman

## Abstract

Mutations in genes that affect mitochondrial function cause primary mitochondrial diseases. Mitochondrial diseases are highly heterogeneous and even patients with the same mitochondrial disease can exhibit broad phenotypic heterogeneity, which is poorly understood. Mutations in subunits of mitochondrial respiratory complex I cause complex I deficiency, which can result in severe neurological symptoms and death in infancy. However, some complex I deficiency patients present with much milder symptoms. The most common nuclear gene mutated in complex I deficiency is the highly conserved core subunit NDUFS1. To model the phenotypic heterogeneity in complex I deficiency we used RNAi lines targeting the *Drosophila* NDUFS1 homolog ND-75 with different efficiencies. Strong knockdown of ND-75 in *Drosophila* neurons resulted in severe behavioural phenotypes, reduced lifespan, altered mitochondrial morphology, reduced endoplasmic reticulum (ER)-mitochondria contacts and activation of the unfolded protein response (UPR). By contrast, weak ND-75 knockdown caused much milder behavioural phenotypes and changes in mitochondrial morphology. Moreover, weak ND-75 did not alter ER-mitochondria contacts or activate the UPR. Weak and strong ND-75 knockdown resulted in overlapping but distinct transcriptional responses in the brain, with weak knockdown specifically affecting proteosome activity and immune response genes. Metabolism was also differentially affected by weak and strong ND-75 knockdown including gamma-aminobutyric acid (GABA) levels, which may contribute to neuronal dysfunction in ND-75 knockdown flies. Several metabolic processes were only affected by strong ND-75 knockdown including the pentose phosphate pathway and the metabolite 2-hydroxyglutarate (2-HG), suggesting 2-HG as a candidate biomarker of severe neurological mitochondrial disease. Thus, our *Drosophila* model provides the means to dissect the mechanisms underlying phenotypic heterogeneity in mitochondrial disease.

## Introduction

Primary mitochondrial diseases are a broad spectrum of disorders characterised by defects in mitochondrial oxidative phosphorylation (OXPHOS). Primary mitochondrial diseases are caused by inherited or spontaneous mitochondrial DNA (mtDNA) or nuclear DNA mutations in genes required for mitochondrial function (Russell et al., 2020). These include genes encoding OXPHOS subunits, enzymes required for mtDNA maintenance, and enzymes involved in regulating mitochondrial gene expression (Gorman et al., 2016).

Mitochondrial diseases can arise both during childhood and adulthood, although this is often dictated by the type of genetic mutation involved. Early-onset mitochondrial diseases, where symptoms develop during infancy or childhood, are often caused by autosomal recessive mutations (Skladal et al., 2003). Adult-onset mitochondrial diseases are largely caused by mtDNA mutations and are generally less severe than those that are early-onset (Gorman et al., 2015). Both early-onset and adult-onset mitochondrial diseases are commonly associated with the development of neurological symptoms, which is likely due to the high metabolic demand of the CNS.

The prevalence of primary mitochondrial diseases is predicted to be ∼ 1 in 7634 at birth, and ∼ 1 in 4300 by adulthood (Skladal et al., 2003; Gorman et al., 2015). As a collective and individually, primary mitochondrial diseases are highly heterogeneous; the age of onset, tissue-specificity, and symptom severity can all vary depending on the individual and gene affected. Mitochondrial diseases are classified based on clinical symptoms, however, the genetic and phenotypic heterogeneity within some of these diseases can make diagnosis very challenging, and often patients do not fit within the defined criteria (Gorman et al., 2016; Finsterer et al., 2018). The underlying reasons for phenotypic heterogeneity within individual primary mitochondrial diseases are not well understood, which makes it challenging to identify common and distinct pathogenic mechanisms. Furthermore, our lack of knowledge surrounding the pathogenesis of mitochondrial diseases has made it difficult to develop therapies.

Complex I deficiency is the most common childhood mitochondrial disease and is caused by mutations in genes encoding complex I structural or assembly components (Fassone and Rahman, 2012). Complex I deficiency encompasses a broad spectrum of disorders with considerable phenotypic heterogeneity (Koene et al., 2012; Bjorkman et al., 2015). NDUFS1 is the most commonly mutated nuclear gene causing complex I deficiency (Fassone and Rahman, 2012). NDUFS1 is an [Fe-S] cluster containing subunit involved in electron transfer that lies within the N module of complex I (Wirth et al., 2016). Mutations in NDUFS1 typically cause severe and rapidly progressive leukoencephalopathy and death within the first two years of life (Kashani et al., 2014). However, phenotypic heterogeneity has been reported in NDUFS1 complex I deficiency, including several paediatric patients with much more mild symptoms (Hoefs et al., 2010; Ferreira et al., 2011; Fassone and Rahman, 2012; Kashani et al., 2014; Bjorkman et al., 2015).

Animal models of mitochondrial disease have been highly successful in revealing underlying mechanisms and identifying potential disease modifying therapies (Duncan and Bateman, 2016; Hunt and Bateman, 2018; Chen et al., 2019; Granat et al., 2020; van de Wal et al., 2022). Mirroring the diversity of mitochondrial diseases, animal models can vary greatly in the range and severity of phenotypes. However, none of these models have recapitulated the phenotypic heterogeneity observed within mitochondrial disease caused by mutations in the same gene. We recently generated a *Drosophila* model of complex I deficiency using knockdown of the complex I subunit NDUFS1 (ND-75) in *Drosophila* (Granat et al., 2023). Knockdown of ND-75 in this model is highly efficient and results in severe neurological symptoms and very early death, reflecting severe complex I deficiency. Here, we have used an independent RNAi against ND-75 that is much less efficient and causes a weaker knockdown. We find that this weak ND-75 RNAi results in far milder behavioural and cellular phenotypes. Importantly, weak knockdown of ND-75 does not cause changes in endoplasmic reticulum (ER)-mitochondria contacts or activation of ATF4 in neurons. Interestingly, weak ND-75 knockdown in neurons causes significant transcriptional and metabolic changes in the brain but these diverge from strong knockdown of ND-75. The genetic tools that we have characterised provide a powerful system for studying the mechanisms contributing to phenotypic heterogeneity in mitochondrial disease.

## Results

### Manipulation of ND-75 knockdown efficiency using RNAi

Mutations in *NDUFS1* cause complex I deficiency and Leigh syndrome (Koene et al., 2012; Lake et al., 2016). NDUFS1 lies within the N module of complex I and is an [Fe-S] cluster containing core subunit involved in electron transfer (Wirth et al., 2016). To model the phenotypic heterogeneity of complex I deficiency we utilised two independent non-overlapping ND-75 RNA interference (RNAi) lines with different knock-down efficiencies, ND-75^KK108222^ and ND-75^HMS00853^, hereafter referred to as ND-75^KDweak^ and ND-75^KDstrong^ respectively (Supplemental Figure S1A).

To compare the effects of ubiquitous ND-75 knockdown using these two RNAi lines in adult *Drosophila* we expressed ND-75^KDweak^ and ND-75^KDstrong^ using *Tub-Gal4*, however this resulted in embryonic lethality. To circumvent this, we used two strategies: 1. The GeneSwitch system (Roman et al., 2001), using *da-GSGAL4* to restrict ubiquitous ND-75 knockdown to adult flies; 2. ubiquitous expression during development using *Tub-GAL4* together with a temperature-sensitive repressor of Gal4, *Tub-GAL80^ts^,* 25°C (Gal80^ts^ is inactive at 30°C and partially active at 25°C (McGuire et al., 2003)).

Ubiquitous expression of ND-75^KDweak^ and ND-75^KDstrong^ in adult flies with *da-GSGAL4* for five days caused a 40% and 88% reduction in ND-75 mRNA levels respectively (Supplemental Figure S1B). Analysis of rotenone-sensitive NADH oxidation in mitochondria from adult flies expressing ND-75^KDweak^ with *da-GSGAL4* showed they had similar complex I activity to controls, while flies expressing ND-75^KDstrong^ had an 87% reduction in complex I activity (Supplemental Figure S1C). Loss of individual complex I subunits can result in collapse of the whole complex (Ugalde et al., 2004). To test this possibility, we analysed the level of ND-30 expression, the orthologue of mammalian NDUFS4 and a component of Q module of complex I (Garcia et al., 2017). ND-30 expression was unaffected in flies expressing ND-75^KDweak^ with *da-GSGAL4* but ND-30 levels were significantly reduced in flies expressing ND-75^KDstrong^ (Supplemental Figure S1D-F). Consistent with these data, ubiquitous expression of ND-75^KDstrong^ but not ND-75^KDweak^ in adult flies using *da-GSGAL4* caused a significant reduction in climbing ability (Supplemental Figure S1G).

ND-75^KDstrong^ expression with *Tub-GAL80^ts^;Tub-GAL4* at 25°C caused developmental lethality. Using *Tub-GAL80^ts^;Tub-GAL4* at 25°C to express ND-75^KDweak^ resulted in a high a degree of pupal lethality and a small number of viable adult escaper flies. These escaper flies had a 79% reduction in ND-75 expression and, although insufficient in number to isolate mitochondria and analyse complex I activity, showed a significant decrease in ND-30 levels and severely reduced climbing ability (Supplemental Figure S1H-L), indicating loss of complex I activity.

Taken together, use of these two ubiquitous expression methods show that loss of complex I activity, the associated loss of ND-30 expression and reduced climbing ability requires at least a 40% reduction in ND-75 gene expression. Moreover, these data demonstrate that both ND-75 RNAi lines are capable of efficient ND-75 knockdown, but ND-75^KDstrong^ is more potent than ND-75^KDweak^.

### Knockdown of ND-75 in neurons models the phenotypic heterogeneity of complex I deficiency

Complex I deficiency primarily affects the nervous system in patients (Koene et al., 2012). To model the phenotypic heterogeneity of complex I deficiency in the nervous system, we induced pan-neuronal expression of ND-75^KDweak^ and ND-75^KDstong^ using *nSyb-GAL4.* ND-75^KDweak^ and ND-75^KDstrong^ expression in neurons using *nSyb-GAL4* caused an inability to climb and pupal lethality respectively (Figure 1A). In the presence of *Tub-Gal80^ts^*at 25°C, pan neuronal expression of ND-75^KDstrong^ using *nSyb-GAL4* resulted in viable flies that were unable to climb, while expression of ND-75^KDweak^ did not cause a climbing phenotype (Figure 1B). We next measured open-field behaviour in pan-neuronal ND-75^KD^ flies. The average speed and total distance moved by ND-75^KDstrong^ flies was dramatically reduced, and immobility was increased compared to control flies (Figure 1C-E). ND-75^KDweak^ flies also had locomotion phenotypes but these were less severe than those observed in ND-75^KDstrong^ flies (Figure 1C-E).

**Figure 1.**
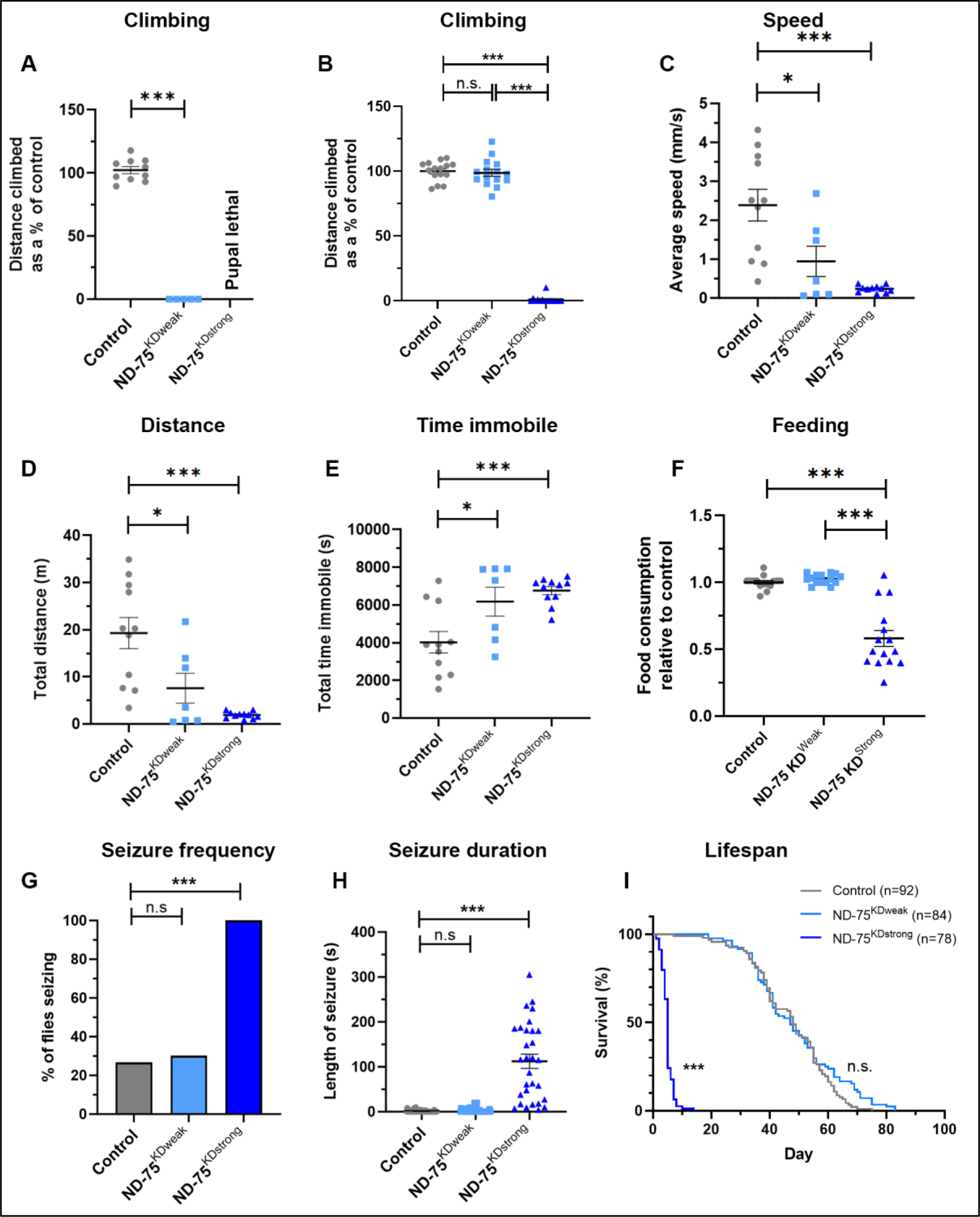
Pan neuronal weak and strong knockdown of ND-75 models the phenotypic heterogeneity in mitochondrial disease. (A) Pan-neuronal expression using *nSyb-Gal4* results in flies that are completely unable to climb with ND-75^KDweak^ and lethality with ND-75^KDstrong^. Control n=10, ND-75^KDweak^ n=5 flies. (B) Pan-neuronal expression using *Gal80^ts^;nSyb-Gal4* at 25°C results in flies that climb normally with ND-75^KDweak^ but are unable to climb with ND-75^KDstrong^. n=15 flies for all genotypes. (C-E) Pan-neuronal expression using *Gal80^ts^;nSyb-Gal4* at 25°C causes mild locomotor phenotypes with ND-75^KDweak^ and strong locomotor phenotypes with ND-75^KDstrong^. Control n=11, ND-75^KDweak^ n=7, ND-75^KDstrong^ n=11 flies. (F) Pan-neuronal expression using *Gal80^ts^;nSyb-Gal4* at 25°C causes reduced feeding with ND-75^KDstrong^ but not ND-75^KDweak^. n=15 flies for all genotypes. (G, H) Pan-neuronal expression using *Gal80^ts^;nSyb-Gal4* at 25°C causes seizures with ND-75^KDstrong^ but not ND-75^KDweak^. Control n=60, ND-75^KDweak^ n=63, ND-75^KDstrong^ n=30 flies. (I) Pan-neuronal expression using *Gal80^ts^;nSyb-Gal4* at 25°C causes greatly reduced lifespan with ND-75^KDstrong^ but not ND-75^KDweak^. Control n=92, ND-75^KDweak^ n=84, ND-75^KDstrong^ n=78 flies. Males flies were used in (A-F, I). Male and female flies were used in (G, H). Controls were *nSyb-Gal4* or *Gal80^ts^;nSyb-Gal4* hemizygotes. Data are represented as mean ± SEM and were analysed using one-way ANOVA with Tukey’s posthoc test, Chi-squared for seizure frequency, or log-rank test for survival curve. n.s not significant, * p<.0.5, ***p < 0.001.

Complex I deficiency patients have problems with feeding (Koene et al., 2012), and so we measured food intake in our *Drosophila* model. Pan-neuronal ND-75^KDstrong^ flies had a strong reduction in food intake, whereas feeding was unaffected in ND-75^KDweak^ flies (Figure 1F).

Alongside motor impairments, seizures are frequently reported in complex I deficiency patients (Finsterer and Zarrouk Mahjoub, 2012; Sofou et al., 2014). Using a mechanical stress-induced seizure assay we found that pan-neuronal expression of ND-75^KDstrong^ in neurons caused a dramatic seizure phenotype, with 94% of flies developing seizures, which were significantly longer than controls and, in some cases, lasting four to five minutes (Figure 1G, H). By contrast expression of ND-75^KDweak^ in neurons did not cause a seizure phenotype (Figure 1G, H).

Complex I deficiency patients typically die within the first few years of life but there is considerable phenotypic heterogeneity (Koene et al., 2012; Bjorkman et al., 2015). Consistent with this, pan-neuronal ND-75^KDstrong^ flies had a dramatically reduced lifespan, with a median survival of five days compared to 48 days for controls. By contrast, the lifespan of ND-75^KDweak^ flies was similar to controls (Fig. 1I).

Overall, these data show that ND-75^KDweak^ and ND-75^KDstrong^ flies exhibit contrasting behavioural and lifespan phenotypes, mirroring the phenotypic heterogeneity of complex I deficiency patients.

### Strong but not weak ND-75 knockdown disrupts ER-mitochondria contacts in neurons

Mitochondrial dysfunction is associated with perturbed mitochondrial morphology both in model systems and patients (Koopman et al., 2005; Ekstrand et al., 2007; Koopman et al., 2007). We therefore used super resolution imaging of mitochondrially-targeted GFP to analyse mitochondrial morphology. Expression of ND-75^KDstrong^ in larval motor neurons using *OK371-Gal4* severely perturbed mitochondrial morphology, causing dramatically increased mitochondrial number and volume (Fig. 2A-E). Expression of ND-75^KDweak^ caused a much less dramatic but still significant increase in mitochondrial number and volume (Figure 2A-E).

**Figure 2.**
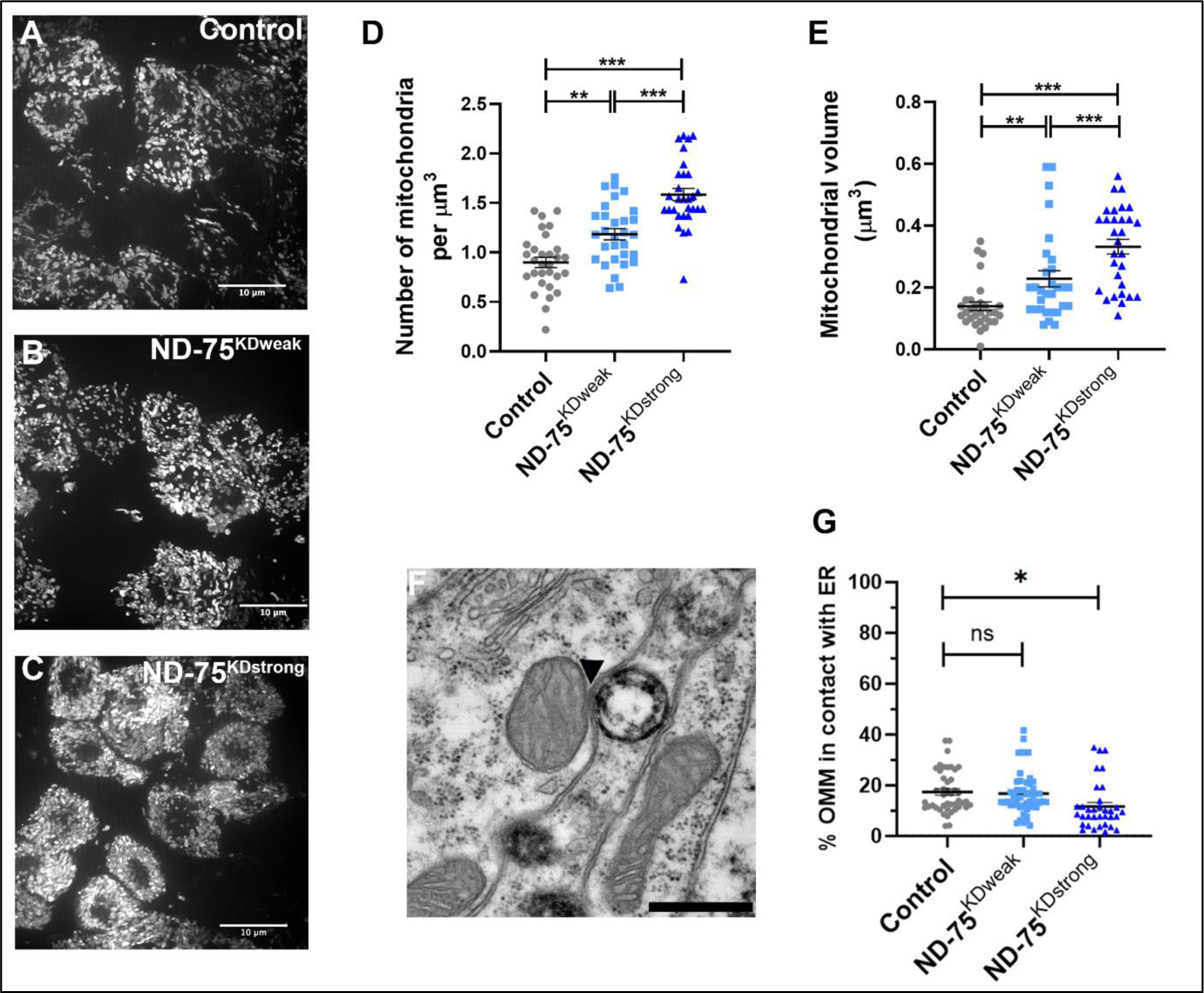
Weak and strong knockdown of ND-75 have different effects on mitochondrial morphology and ER-mitochondria contacts. (A-C) Expression of mitochondria-targeted GFP to visualise mitochondria in control (A), ND-75^KDweak^ (B) and ND-75^KDstrong^ (C) larval motor neurons using *OK371-Gal4*. Images were taken using iSIM. (D, E) Quantification of mitochondrial number and volume in larval motor neurons with ND-75 knockdown using *OK371-Gal4*. Control n=30, ND-75^KDweak^ n=30, ND-75^KDstrong^ n=30 ROIs. (F) Example transmission electron microscopy image of mitochondria in the adult brain. Arrowhead indicates ER-mitochondria contact. (G) Quantification of ER-mitochondria contacts in adult brain from control or with pan-neuronal ND-75 knockdown (using *Gal80^ts^;nSyb-Gal4*). Control n=44, ND-75^KDweak^ n=49, ND-75^KDstrong^ n=34 mitochondria. Controls were *OK371-Gal4* or *Gal80^ts^;nSyb-Gal4* hemizygotes. Data are represented as mean ± SEM and were analysed using one-way ANOVA with Tukey’s posthoc test. n.s not significant, * p<.0.5, **p<0.01, ***p < 0.001.

Mitochondria and the endoplasmic reticulum (ER) are frequently apposed and connected through ER-mitochondria contacts, which can be observed at the ultrastructural level (Figure 2F) (Stoica et al., 2014). We therefore used transmission electron microscopy to visualise and quantify ER-mitochondria contacts in the adult brain. Pan-neuronal ND-75^KDstrong^ expression using *Tub-Gal80^ts^;nSyb-Gal4* caused a decrease in the number of ER-mitochondria contacts, but ER-mitochondria contacts in pan-neuronal ND-75^KDweak^ flies were similar to control (Figure 2H). These data suggest that exceeding a threshold of complex I inactivation is required to perturb ER-mitochondria contacts in neurons.

### The ER EPR is activated by strong but not weak ND-75 knockdown in neurons

Disruption of ER homeostasis leads to ER stress and activation of the unfolded protein response (UPR) (Oslowski and Urano, 2011). The UPR has been consistently shown to be activated by mitochondrial dysfunction (Hunt et al., 2019; Granat et al., 2020). To investigate whether ND-75 knockdown triggers UPR activation in neurons, we used activating transcription factor 4 (ATF4) expression as a readout of UPR activation. Expression of ND-75^KDstrong^ but not ND-75^KDweak^ using *OK371-Gal4* in larval neurons or *Tub-Gal80^ts^;nSyb-Gal4* in the adult brain caused strong activation of ATF4 expression (Figure 3A-F). Triggering of the UPR activates protein kinase R-like endoplasmic reticulum kinase (PERK), resulting in ATF4 up-regulation through the phosphorylation of the translation initiation factor eIF2α (Hetz and Mollereau, 2014). Consistent with this, ND-75^KDstrong^ but not ND-75^KDweak^ expression increased eIF2α levels in neurons (Figure 3G-J). Therefore, efficient knockdown of ND-75 activates the UPR and ATF4 in neurons.

**Figure 3.**
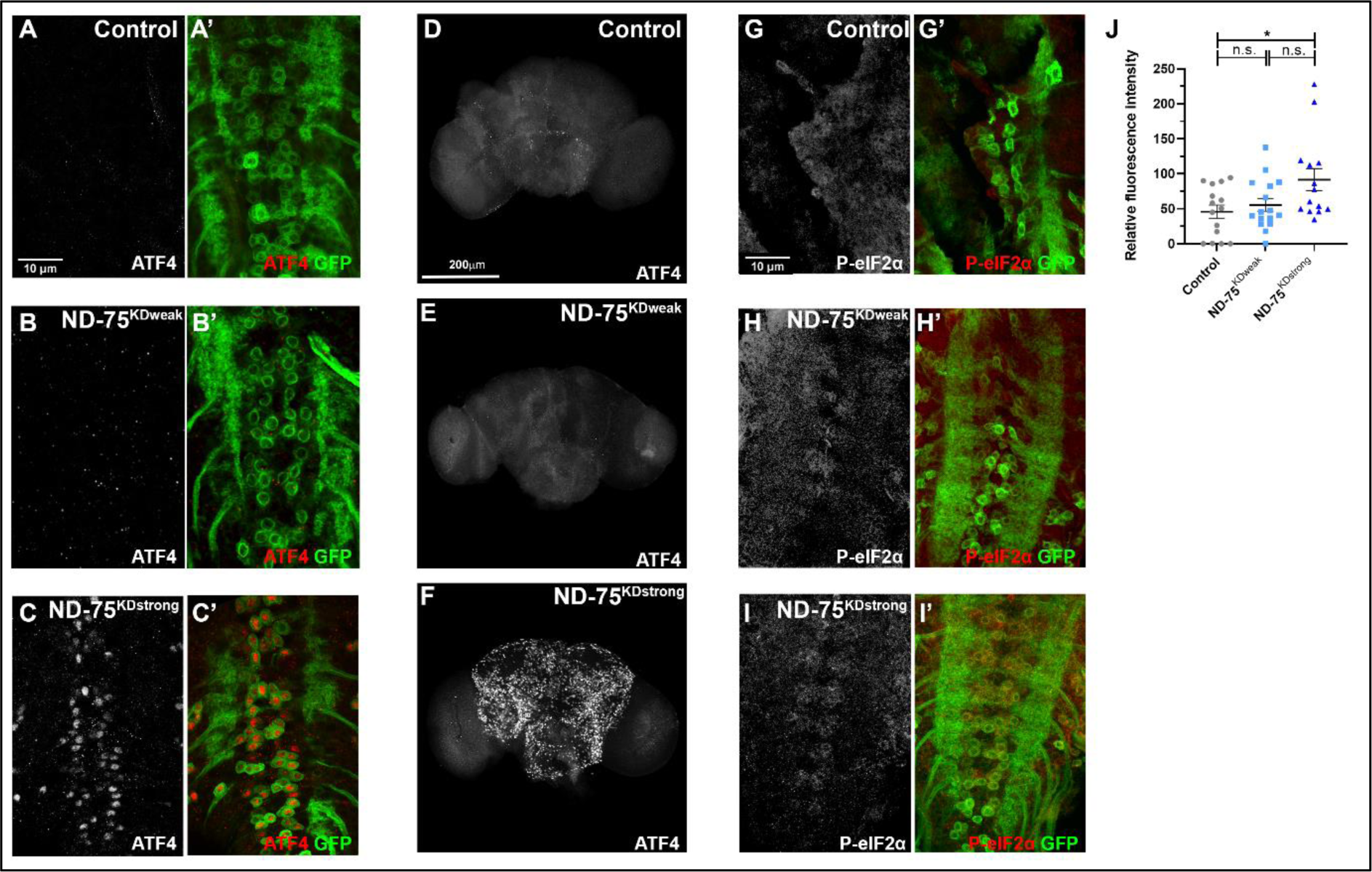
Strong but not weak ND-75 knockdown activates the ER UPR. (A-C). ATF4 (red) expression in control (A), ND-75^KDweak^ (B) and ND-75^KDstrong^ (C) larval motor neurons using *OK371-Gal4*. CD8-GFP (green) expression labels motor neurons. (D-F) ATF4 expression in control (D), ND-75^KDweak^ (E) and ND-75^KDstrong^ (F) adult brain tissue using *Tub-Gal80^ts^*; *nSyb-Gal4*. 1 day old flies were used. (G-I) Phospho-eIF2α (P-eIF2α, red) expression in control (G), ND-75^KDweak^ (H) and ND-75^KDstrong^ (I) larval motor neurons using *OK371-Gal4*. CD8-GFP (green) expression labels motor neurons. (J) Quantification of Phospho-eIF2α expression. Control n=15, ND-75^KDweak^ n=16, ND-75^KDstrong^ n=14 larval CNS. Data were analysed using ANOVA with Tukey’s posthoc test. Controls were *OK371-Gal4* or *Tub-Gal80^ts^*; *nSyb-Gal4* hemizygotes. Data are represented as mean ± SEM. n.s. not significant, *p<0.05.

### Weak and strong ND-75 knockdown have different effects on transcription and metabolism in the brain

The activation of ATF4 indicated that strong ND-75 knockdown would result in changes to the transcriptome in neurons. In order to understand the differences in the transcriptional response to pan-neuronal weak and strong ND-75 knockdown using *Tub-Gal80^ts^;nSyb-Gal4*, we performed RNA sequencing of fly heads. Principle component analysis (PCA) showed the two ND-75 knockdown groups were well separated from each other and the control (Figure 4A). In total there were 1127 differentially expressed genes (DEGs) in ND-75^KDweak^ flies compared to control, of which 501 were up- and 626 genes were downregulated (Figure 4B, C; Supplemental tables S1, S2). Surprisingly, ND-75^strong^ flies showed a lower number of 508 DEGs (Figure 4B, D; Supplemental tables S3, S4). Of these, 373 were upregulated, while 135 were downregulated (Figure 4B, D; Supplemental tables S3, S4). Although there were fewer DEGs, the average fold change of the upregulated DEGs in ND-75^KDstrong^ flies (FC=8.41) was significantly higher than in ND-75^KDweak^ flies (FC=5.2, p=1.56×10^-5^), indicating that strong ND-75 knockdown elicits a more pronounced increase in DEG expression. Strikingly, the first and fourth most significantly increased DEGs in ND-75^KDstrong^ flies were *MFS3* and *Ldh*, encoding the glucose/trehalose transporter Major Facilitator Superfamily Transporter 3 and lactate dehydrogenase, respectively (Figure 4D; Supplemental table S3). By contrast, neither *MFS4* nor *Ldh* were differentially expressed in ND-75^KDweak^ flies. Increased *MFS3* and *Ldh* expression indicate a pronounced shift to glycolytic metabolism as a result of strong CI deficiency and activation of ATF4, which regulates *Ldh* expression in *Drosophila* (Lee et al., 2015; Hunt et al., 2019).

**Figure 4.**
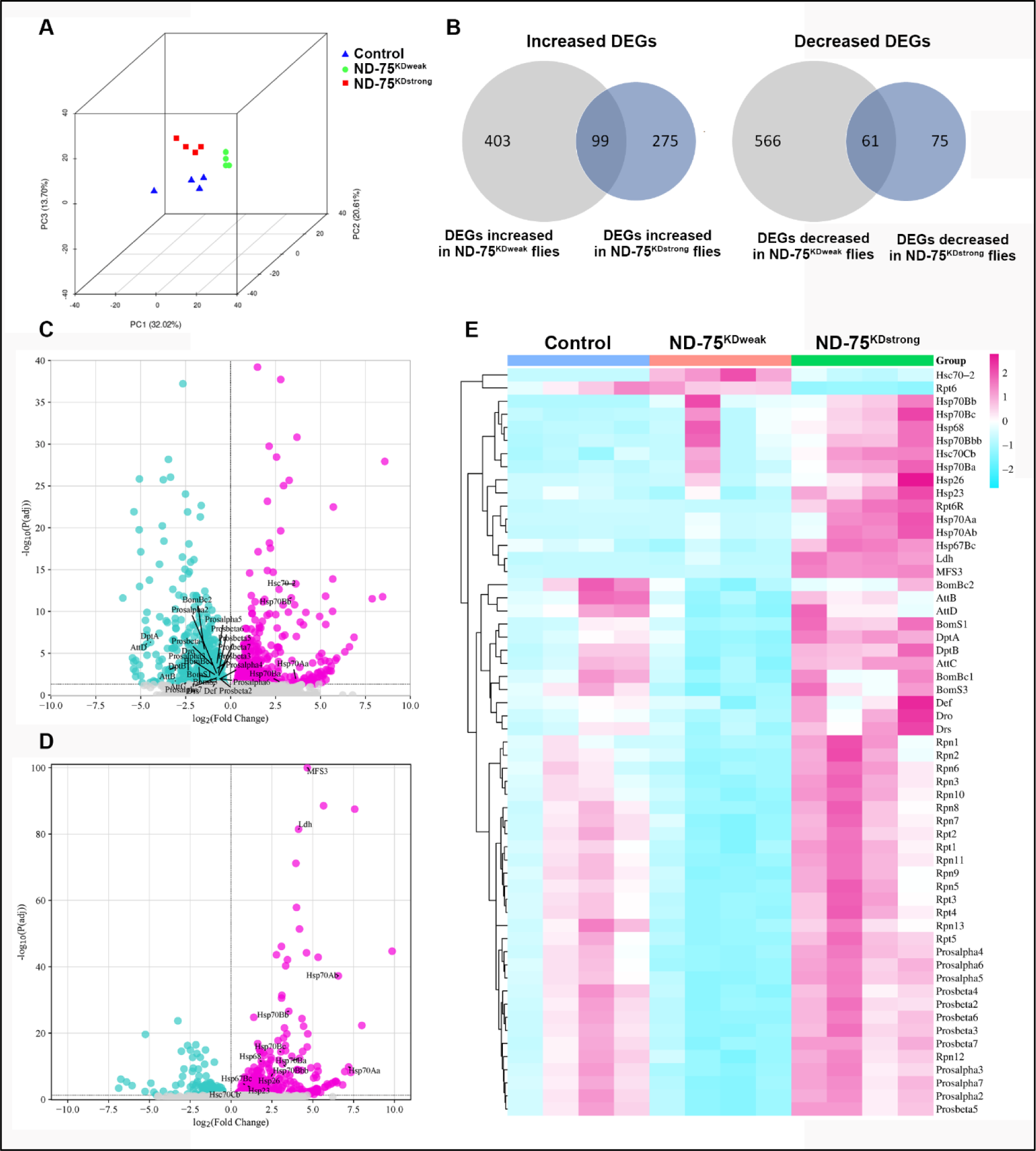
Weak and strong ND-75 knockdown cause overlapping but distinct transcriptome changes. (A) PCA plot of transcriptome analysis from head tissue of control, and flies expressing ND-75^KDweak^ and ND-75^KDstrong^ in neurons using *Tub-Gal80^ts^*; *nSyb-Gal4*. (B) Numbers of DEGs increased and decreased in ND-75^KDweak^ and ND-75^KDstrong^ flies and their overlap. (C, D) Volcano plots of DEGs in ND-75^KDweak^ (C) and ND-75^KDstrong^ (D) flies. (E) Heat map showing relative expression of representative genes in ND-75^KDweak^ and ND-75^KDstrong^ flies. 2 day old male and female flies were used. Controls were *Tub-Gal80^ts^*; *nSyb-Gal4* hemizygotes.

Gene ontology (GO) analysis showed ND-75^KDweak^ upregulated DEGs were significantly enriched for genes involved in protein misfolding, oxidoreductase activities, heme/iron binding and lipid catabolic processes, while downregulated DEGs were enriched for proteasome activity and immune response genes (Figure 4C, E; Supplemental Figure S2A, B). GO analysis of upregulated DEGs in ND-75^KDstrong^ highlighted the unfolded protein response, oxidoreductase activities, lipid catabolism and heme binding (Supplemental Figure S3C), while no GO categories were significantly downregulated in ND-75^KDstrong^ DEGs.

The molecular function ‘misfolded protein binding’ was enriched in upregulated DEGs in both conditions, however, although the total number of DEGs in ND-75^KDweak^ flies was more than double ND-75^KDstrong^ flies, expression of 11 heatshock genes (*Hsp23, Hsp26, Hsp67Bc, Hsp68, Hsp70Aa, Hsp70Ab, Hsp70Ba, Hsp70Bb, Hsp70Bbb, Hsp70Bc*) were significantly increased in ND-75^KDstrong^ flies while only 4 (*Hsc70-2, Hsp70Aa, Hsp70Ba, Hsp70Bb*) were increased in ND-75^KDweak^ flies (Figure 4D, E; Supplemental tables S1, S3). These data suggest that strong CI deficiency elicits a greater chaperone response than weak CI deficiency in the brain.

Subunits of the 26S proteasome (e.g. *Prosalpha4, Prosalpha5, Prosbeta7,*) were strongly enriched in the ND-75^KDweak^ downregulated DEGs, whereas no 26S proteasome subunits were downregulated in ND-75^KDstrong^ flies (Figure 4C-E; Supplemental tables S2, S4). The expression of many different antimicrobial peptides (e.g. *AttB*, *AttD*, *DptA,*) were downregulated in ND-75^KDweak^ flies, whereas only one (*Drsl4*) was downregulated in ND-75^KDstrong^ flies (Figure 4C-E; Supplemental tables S2, S4). Therefore, weak CI deficiency specifically represses proteosome activity and immune response genes in the brain.

Complex I deficiency causes profound alterations in brain metabolism (McElroy et al., 2020; van de Wal et al., 2022; Granat et al., 2023). In keeping with this, our transcriptomic analysis showed ND-75 knockdown results in altered expression of metabolic genes such as *Ldh*. To understand how weak and strong complex I deficiency in *Drosophila* neurons affects metabolism, we performed untargeted metabolomics of head tissue from flies expressing ND-75^KDweak^ and ND-75^KDstrong^ in neurons using *Gal80^ts^;nSyb-Gal4.* PCA revealed that knockdown of ND-75 using either ND-75^KDweak^ or ND-75^KDstrong^ produced metabolic profiles that were distinct from control (Figure 5A). The profiles were also distinct from each other, with ND-75^KDweak^ bearing more similarity to controls than ND-75^KDstrong^ (Figure 5A).

**Figure 5.**
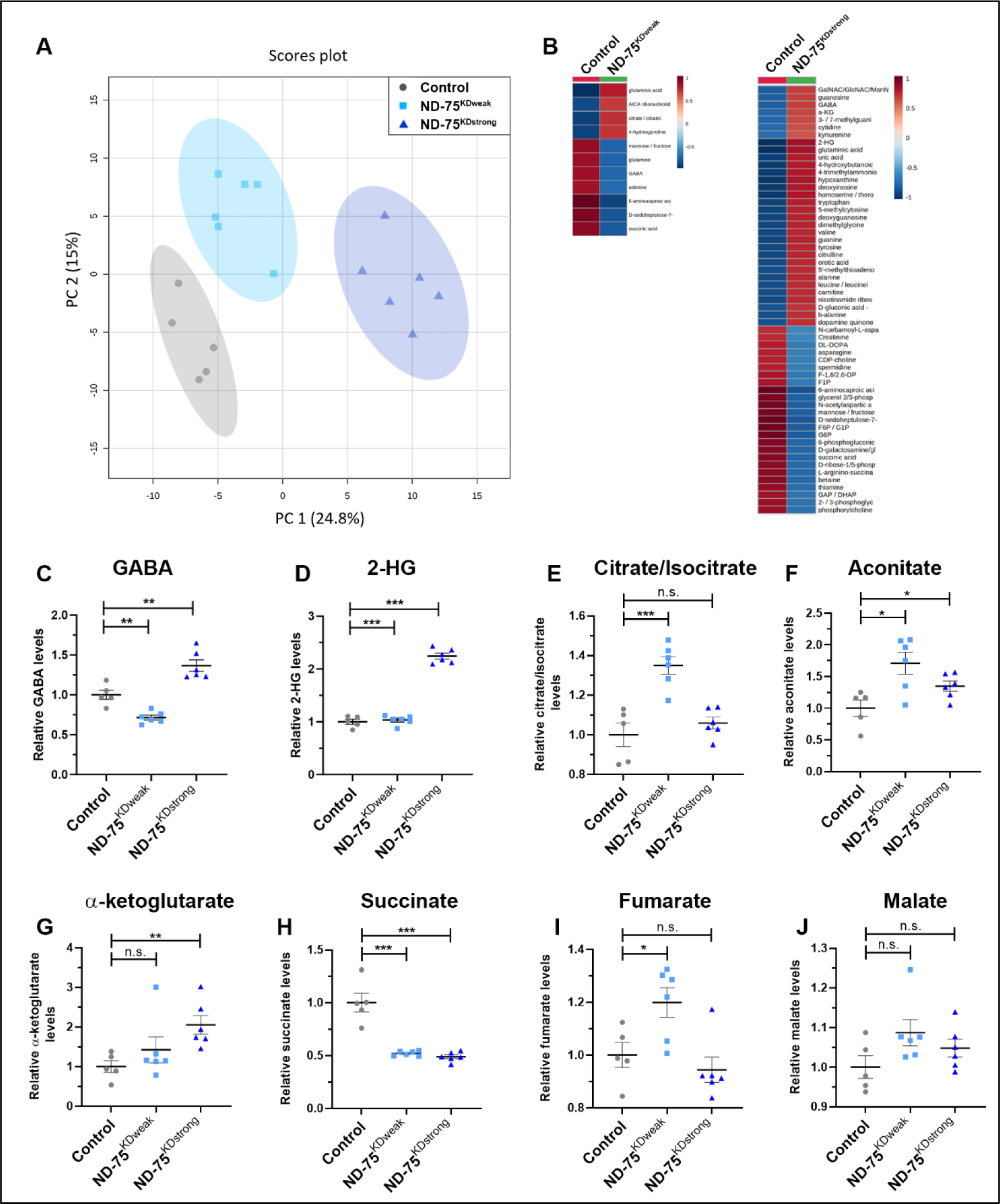
Metabolic reprogramming in neurons is dependent on the level of ND-75 knockdown. (A) PCA plot of metabolomic analysis from head tissue of control, and flies expressing ND-75^KDweak^ and ND-75^KDstrong^ in neurons using *Tub-Gal80^ts^*; *nSyb-Gal4*. (B) Metabolites significantly misregulated in ND-75^KDweak^ and ND-75^KDstrong^ flies. Scale represetnts the z score. (C, D) Levels of GABA (C) and 2-HG (D) in ND-75^KDweak^ and ND-75^KDstrong^ flies. (E-J) Levels of TCA cycle metabolites in ND-75^KDweak^ and ND-75^KDstrong^ flies. 2 day old male and female flies were used. Control n=5, ND-75^KDweak^ n=6, ND-75^KDstrong^ n=6 biological replicates. Data were analysed using ANOVA with Tukey’s posthoc test. Controls were *Tub-Gal80^ts^*; *nSyb-Gal4* hemizygotes. Data are represented as mean ± SEM. n.s. not significant, *p<0.05, **p<0.01, ***p < 0.001.

Of the 195 metabolites identified (Supplemental Table S5), expression of ND-75^KDweak^ led to a significant up-regulation in the levels of 4 metabolites, and a significant down-regulation in the levels of 7 metabolites compared to controls (Figure 5B, Supplemental Table S6). Expression of ND-75^KDstrong^ led to a significant up-regulation in the levels of 32 metabolites, and a significant down-regulation in the levels of 26 metabolites compared to controls (Figure 5B, Supplemental Table S7). Six of the 11 metabolites mis-regulated in ND-75^KDweak^ flies were also mis-regulated in ND-75^KDstrong^ flies. Interestingly, five of these were similarly increased or decreased in both conditions (6-aminocaproic acid, D-sedoheptulose-7-phosphate, glutaminic acid, succinic acid and mannose/fructose), whereas gamma-aminobutyric acid (GABA) levels were decreased in ND-75^KDweak^ flies but increased in ND-75^KDstrong^ flies (Figure 5C; Supplemental Figure 3A, B; Supplemental Tables S6, S7). As the major inhibitory neurotransmitter, GABA may contribute to the behavioural defects caused by strong ND-75 knockdown.

We performed metabolite set enrichment analysis (MSEA) to identify biological processes or molecular pathways that were significantly dysregulated (Xia and Wishart, 2010). ND-75^KDweak^ expression led to significant dysregulation in 22 of 54 (41%) human KEGG pathways (Supplemental Figure S3C). In comparison, ND-75^KDstrong^ expression caused significant dysregulation in 44 of 54 KEGG pathways examined (81%) (Supplemental Figure S3D). Comparison of the weak and strong ND-75 knockdown MSEA highlighted several processes that were dysregulated specifically in ND-75^KDstrong^ flies including the pentose phosphate pathway, pantothenate and CoA biosynthesis, beta alanine metabolism, lysine degradation and galactose metabolism (Supplemental Figure S3C, D). Moreover, 2-hydroxyglutarate (2-HG) levels were strongly increased in ND-75^KDstrong^ but not ND-75^KDweak^ flies (Figure 5D). 2-HG is synthesised from the TCA cycle intermediate α-ketoglutarate and we and others have shown 2-HG levels are increased as a consequence of mitochondrial dysfunction and contribute to neuronal dysfunction (Burr et al., 2016; Hunt et al., 2019).

MSEA revealed that the TCA cycle was dysregulated in both ND-75^KDweak^ and ND-75^KDstrong^ flies (Supplemental Figure S3C, D). Mitochondrial complex I is functionally coupled to the TCA cycle. NADH oxidation by FMN within the matrix arm of complex I replenishes NAD+, a co-factor required for the generation of α-ketoglutarate from isocitrate, succinyl-CoA from α-ketoglutarate, and oxaloacetate from malate (Supplemental Figure S3E). Expression of ND-75^KDweak^ led to a significant rise in citrate/isocitrate levels (Figure 5E), whereas expression of ND-75^KDstrong^ had no significant effect (Figure 5E). In contrast, both the ND-75^KDweak^ and ND-75^KDstrong^ induced a significant increase in aconitate, an intermediate in the conversion of citrate to isocitrate (Figure 5F). Consistent with the changes in 2-HG levels, expression of ND-75^KDstrong^ but not ND-75^KDweak^ caused a significant elevation in α-ketoglutarate levels (Figure 5G), and both produced a significant, comparable reduction in succinate (succinic acid) levels (Figure 5H). Expression of ND-75^KDweak^, but not ND-75^KDstrong^ led to a significant increase in fumarate levels (Figure 5I), whereas neither had any significant effect on malate levels (Figure 5J). Succinyl-CoA and oxaloacetate, the remaining TCA cycle intermediates, were not detected. Taken together, these data demonstrate that neuronal-specific knockdown of ND-75 leads to significant metabolic abnormalities in a range of metabolic pathways, including the TCA cycle, and that the severity of metabolic dysfunction is dependent on the extent of ND-75 knockdown.

## Discussion

Primary mitochondrial diseases have a common cause but are highly heterogeneous. We have modelled this heterogeneity using a knockdown approach in *Drosophila*. Using two independent ND-75 RNAi lines, our data show that the behavioural phenotypes and mitochondrial morphology defects caused by complex I deficiency in neurons correlate with the ND-75 knockdown efficiency. However, the loss of ER-mitochondria contacts and activation of the UPR only occur with strong ND-75 knockdown. The transcriptional and metabolic changes caused by weak and strong ND-75 knockdown provide mechanistic insight and highlight specific molecular functions and metabolic pathways that differentiate the heterogeneity in complex I deficiency. Thus, our *Drosophila* model has good face validity for complex I deficiency and provides new insight into the cellular and molecular mechanisms associated with phenotypic heterogeneity.

Complex I deficiency typically manifests with neonatal-onset lactic acidosis or encephalomyopathies, Leigh syndrome, leukoencephalopathy, hypertrophic cardiomyopathy, exercise intolerance and is often fatal (Fassone and Rahman, 2012). However, heterogeneity has been documented in patients with mutations in several different complex I subunits. Missense mutations in the assembly factor NDUFAF5 cause low complex I activity resulting in classical early onset Leigh syndrome with onset before six months and death by three years (Gerards et al., 2010). However, several adult patients have been reported with more mild loss of complex I activity and minor neurological involvement (Gerards et al., 2010; Simon et al., 2019). Moreover, paediatric patients with mutations in the complex I accessory subunit NDUFA12 exhibited a range in the onset of motor symptoms (Torraco et al., 2021). Patients with NDUFS1 mutations most frequently manifest with severe neonatal rapidly progressive leukoencephalopathy that is fatal. However, a seven-year-old patient with a missense mutation in NDUFS1 had developmental delay and early motor symptoms associated with infections that gradually improved to the point where he was in generally good health (Kashani et al., 2014). This patient’s fibroblasts showed complex I activity was reduced to 20% of control cells. Björkman et al., described three patients with NDUFS1 mutations, two with severe symptoms who died within six weeks of birth and a third three-and-a-half-year-old with much milder symptoms, including hypertonia, who had normal language development and crawling (Bjorkman et al., 2015). Pyruvate + malate and NADH ferricyanide reductase activities in isolated skeletal muscle mitochondria were outside the normal range but more mildly affected in this patient than the severe cases (Bjorkman et al., 2015). Although limited in number, these clinical studies demonstrate that the mild behavioural phenotypes in ND-75^KDweak^ flies reflect the symptoms caused by rare NDUFS1 mutations that likely cause a less severe loss of complex I activity.

The two ND-75 RNAi lines used both target non-overlapping regions towards the end of the gene (Supplemental Figure S1A). The difference in efficiency between the two RNAis can likely be explained by their properties; ND-75^KDstrong^ encodes a short hairpin RNA (shRNA), whereas ND-75^KDweak^, encodes a long hairpin RNA (lhRNA) (Supplemental Figure S1A). Previous comparisons of shRNAs and lhRNAs targeting same gene have revealed that shRNAs consistently produce stronger phenotypes than lhRNAs (Ni et al., 2011; Bartoletti et al., 2017), strongly suggesting that shRNAs, which mimic endogenous microRNAs, are more efficient at reducing mRNA levels. Consistent with this, ubiquitous knockdown (using *Da-Gal4*) of ND-75 with an independent lhRNA produced viable flies with an approximately 50% reduction in lifespan (Hegde et al., 2014). It will be interesting in future to employ the RNAi lines we have characterised to model the effects of complex I deficiency and phenotypic heterogeneity in other tissues, such as muscle, affected in mitochondrial disease patients.

ND-75^KDstrong^ flies had loss of climbing ability, severe loss of locomotor function, seizures and greatly reduced lifespan. By contrast, the only behavioural defect in ND-75^KDweak^ flies was a mild reduction in locomotor function. However, several cellular and molecular changes were evident in ND-75^KDweak^ flies, including increased mitochondrial number and volume in neurons, 1127 DEGs and altered levels of 11 metabolites in the brain. Significantly, ND-75^KDweak^ expression did not activate ATF4, suggesting that the severe behavioural phenotypes caused by strong ND-75 knockdown are associated with ATF4 expression in neurons. Consistent with this, expression of *Ldh*, which is an ATF4 target in *Drosophila* (Lee et al., 2015; Hunt et al., 2019), is strongly increased in ND-75^KDstrong^ but not in ND-75^KDweak^ flies. Moreover, we previously showed in an independent *Drosophila* mitochondrial dysfunction model that activation of ATF4 in neurons causes increased 2-HG levels in the brain (Hunt et al., 2019). 2-HG is also increased in cells from patients with a form of Leigh syndrome (Burr et al., 2016). Interestingly, metabolomic analysis of children with mutations in respiratory chain complex I, complex III or multiple complex deficiencies showed they had significantly increased 2-HG levels in urine (Reinecke et al., 2012). Thus, 2-HG may have potential as a prognostic biomarker for severe mitochondrial disease involving activation of ATF4.

The transcriptomic and metabolomic data also show that large scale transcriptional and limited metabolic changes in the brain of ND-75^KDweak^ flies occur in the absence of ATF4 activation. These transcriptional and metabolic changes indicate distinct (e.g. proteosome activity and immune response genes) responses to mild complex I deficiency that do not involve UPR activation. It will be interesting in future to investigate the signalling pathways and transcription factors responsible.

Overall, our study illustrates how using RNAis with different efficiencies to knockdown the same OXPHOS subunit in *Drosophila* provides a powerful means of modelling mitochondrial disease phenotypic heterogeneity. It will be interesting to compare these findings with future studies in complex I deficiency patients and of patient-derived cells to determine the predictive power of our *Drosophila* model.

## Materials and Methods

### Fly strains and growth conditions

Flies were maintained on standard food (per litre: 6.4 g Agar (Fisher), 64 g glucose (Sigma), 16 g ground yellow corn and 80 g Brewer’s yeast (MP Biomed Europe), 3 ml propionic acid (Fisher), 1.8 g methyl 4-hydroxybenzoate (Sigma), 16 ml ethanol (Sigma)) at 25°C in a 12 hour light/dark cycle unless stated otherwise. Genotypes for all experiments are described in Supplemental Table S8. ND-75^KDweak^ (ND-75^KK108222^) was from the Vienna *Drosophila* Resource Center (Dietzl et al., 2007), ND-75^KDstrong^ (ND-75^HMS00853^) was from the TRiP collection (Perkins et al., 2015) and obtained from the Bloomington Stock Center. *Daughterless-GeneSwitch-GAL4* and *Tub-Gal80^ts^; nSyb-Gal4* expression were performed as in (Granat et al., 2023).

### Behavioural analysis

Climbing assays were performed using male flies as in (Hunt et al., 2019). For behavioural assays with ND-75^KDstrong^ flies, vials were placed on their sides during eclosion to prevent flies from becoming stuck in the food. Open-field locomotor activity, seizure, lifespan and CApillary FEeder assay (CAFE) analysis was performed as in (Granat et al., 2023).

### qRT-PCR

qRT-PCR was performed as in (Granat et al., 2023).

### Mitochondrial NADH dehydrogenase (complex I) assay

Complex I assay was performed as in (Granat et al., 2023)

### Immunofluorescence and imaging

Images were taken using a Nikon A1R confocal microscope or a Nikon Vt-iSIM super resolution microscope with NIS Elements software. Confocal imaging and quantification was performed as in (Granat et al., 2023). A Vt-iSIM super resolution microscope (Nikon) was used to capture mitoGFP expression as in (Granat et al., 2023). Primary antibodies were rat anti-ATF4 (1:200, (Hunt et al., 2019)) and rabbit anti-P-eIF2α (1:500, anti-Phospho-eIF2α [Ser51], Cell Signaling Technology 9721). Secondary antibodies were goat anti-rabbit Alexa Fluor 546 (Invitrogen A11035) and goat anti-rat Alexa Fluor 555 (Thermo Fisher A21434).

### Western blot analysis

Western blot analysis was performed as in (Granat et al., 2023). Antibodies used were rabbit anti-VDAC (1:1000; ab14374, Abcam), mouse anti-Ndufs3 (1:500; ab14711, Abcam) and rabbit anti-actin (1:5000; 4967, Cell Signalling Technology).

### RNA sequencing (RNA-Seq) transcriptomic analysis

20 snap frozen fly heads (10 male and 10 female, 2 days old) were used for each replicate and placed into 100 µL of lysis buffer + β-mercaptoethanol from the Absolutely RNA Microprep kit (Agilent Technologies). Each genotype was prepared in quadruplicate. RNA was extracted from using the Absolutely RNA Microprep kit according to the manufacturer’s protocol. The samples were sent on dry ice to Novogene Ltd. Sequencing libraries were generated using NEBNext Ultra TM RNA Library Prep Kit for Illumina (NEB, USA) following manufacturer’s recommendations. RNA-seq was performed as described previously (Granat et al., 2023). Genes with an adjusted P value < 0.05 were assigned as differentially expressed. GO enrichment was performed using the DAVID Knowledgebase (https://david.ncifcrf.gov/tools.jsp). Heatmaps and volcano plots were generated using SRPlot (Tang et al., 2023). RNA-Seq data have been deposited in NCBI’s Gene Expression Omnibus (Edgar et al., 2002) and are accessible through GEO Series accession number GSE248363.

### Metabolomic analysis

Metabolomics was performed as in (Granat et al., 2023). Metabolomic data were analysed using MetaboAnalyst 5.0 (Chong et al., 2018).

### Transmission electron microscopy (TEM)

TEM of adult brain was performed as described previously (Cagin et al., 2015; Granat et al., 2023).

### Statistical analyses

Continuous data are expressed as mean ± S.E.M unless stated otherwise. Non-continuous data are expressed as percentages unless stated otherwise. All data apart from transcriptomic and metabolomic were analysed using Prism 8 (GraphPad). Student’s unpaired two-way t-tests were used for pairwise comparisons of continuous data. An F-test was used to test for unequal variances, and where significant, Welch’s correction was applied to the t-test. A one-way ANOVA with Tukey’s post-hoc test was used for continuous data with multiple comparisons. Chi-squared test and Fisher’s tests were used for non-continuous data, and were applied to the raw values rather than percentages. The log-rank test was used for lifespan data. For the NADH dehydrogenase activity assay data were normalised to the control and then transformed using log base 2. P values <0.05 were considered significant; * p<0.05, ** p<0.01, *** p<0.001.

## Acknowledgements

Stocks obtained from the Bloomington *Drosophila* Stock Center (NIH P40OD018537) and the Vienna *Drosophila* Resource Center were used in this study. We are grateful to the King’s College London Centre for Ultrastructural Imaging for technical assistance and the Wohl Cellular Imaging Centre for help with light microscopy. This work was funded by Alzheimer’s Research UK (ARUK-IRG2017A-2) and the MRC (MR/V013130/1) to JMB; LG was supported by the UK Medical Research Council (MR/N013700/1) and King’s College London MRC Doctoral Training Partnership in Biomedical Sciences; R.P.C was supported by a Northwestern University Pulmonary and Critical Care Department Cugell predoctoral fellowship.

